# Moss Kinesin-14 KCBP accelerates chromatid motility in anaphase

**DOI:** 10.1101/629733

**Authors:** Mari Yoshida, Moé Yamada, Gohta Goshima

## Abstract

KCBP is a microtubule (MT) minus-end-directed kinesin widely conserved in plants. It was shown in *Arabidopsis* that KCBP controls trichome cell shape by orchestrating MT and actin cytoskeletons using its tail and motor domains. In contrast, the *KCBP* knockout (KO) line in the moss *Physcomitrella patens* showed a defect in nuclear and organelle positioning in apical stem cells. Moss KCBP is postulated to transport the nucleus and chloroplast via direct binding to their membranes, since it binds to and transports liposomes composed of phospholipids *in vitro*. However, domains required for cargo transport *in vivo* have not been mapped. Here, we performed a structure-function analysis of moss KCBP. We found that the FERM domain in the tail region, which is known to bind to lipids as well as other proteins, is essential for both nuclear and chloroplast positioning, whereas the proximal MyTH4 domain plays a supporting role in chloroplast transport. After anaphase but prior to nuclear envelope re-formation, KCBP accumulates on the chromosomes, in particular at the centromeric region in a FERM-dependent manner. In the *KCBP* knockout line, poleward chromosome motility in anaphase was reduced and lagging chromosomes occasionally appeared. These results suggest that KCBP binds to non-membranous naked chromosomes via an unidentified protein(s) for their transport. Finally, the liverwort orthologue of KCBP rescued the chromosome/chloroplast mis-positioning of the moss *KCBP* KO line, suggesting that the cargo transport function is conserved at least in bryophytes.

## Introduction

Microtubule (MT) and actin cytoskeletons function as tracks for various intracellular cargos during transportation, such as large organelles in the cytoplasm. KCBP is a MT minus-end-directed kinesin-14 family protein uniquely evolved in the plant lineage (Hamada, 2007; Reddy et al., 1996). In the moss *Physcomitrella patens*, KCBP has been shown to be necessary for proper positioning of the nucleus and chloroplasts (Yamada et al., 2017). Several data led us to propose a model in which KCBP directly binds to nuclear and chloroplast membranes, walks along the MT towards minus ends, and transports those organelles to their proper positions. First, intracellular motility of these organelles is MT-dependent, and not actin-dependent (Miki et al., 2015). Second, the motility of the nucleus and chloroplasts is skewed in the absence of KCBP (Yamada et al., 2017). Overall MT polarity in the observed knockout (KO) cell supports the notion that MT minus-end-directed motility is specifically suppressed without KCBP. Third, recombinant KCBP binds to and transports liposomes composed of acidic phospholipids towards MT minus ends *in vitro* (Jonsson et al., 2015; Yamada et al., 2017). Fourth, KCBP is enriched at the nuclear region during translocation, which takes place at the mitotic telophase in the cell cycle (Yamada et al., 2017). However, more data is necessary to support (or revise) the model.

An open question is whether or not the lipid binding of KCBP drives organelle transport. Although KCBP efficiently binds to liposomes *in vitro*, its enrichment around the chromosomal region is limited to ~20 min after anaphase chromatid separation in the cell; during interphase, no KCBP localisation is observed at the nuclear envelope (NE) and the *KCBP* KO line displays no nuclear migration defect (Yamada et al., 2017). Instead, KCH kinesin was identified as the minus-end-directed transporter of the nucleus during interphase (Yamada and Goshima, 2018). The ‘nuclear’ enrichment of KCBP after chromosome segregation was concluded based solely on its co-visualisation with histone-RFP, which is a chromosomal marker; therefore, it remains unknown if the NE has been re-formed at the time of KCBP localisation and function. Furthermore, nuclear migration is mediated by the NE-embedded LINC (Linker of Nucleoskeleton and Cytoskeleton) or LINC-like complex in other animal, plant and yeast systems; LINC binds to the tail region of motors (dynein, kinesin, myosin) or directly to actin filaments, thereby linking the nucleus to cytoskeleton (Chang et al., 2015; Gundersen and Worman, 2013; Tamura et al., 2013). Second, it remains to be tested whether KCBP drives nuclear and chloroplast translocation independently. There remains a possibility that mis-localised chloroplasts can affect nuclear distribution and vice versa. Recently, it was suggested that abnormal vacuole distribution prevents nuclear translocation in *Arabidopsis* embryos, despite the presence of intact motors and cytoskeleton (Kimata et al., 2019). Third, distinct biochemical activities and cellular functions have been reported for *Arabidopsis thaliana* KCBP. The non-motor domain of AtKCBP binds to actin and MTs, through which it orchestrates MTs and the actin cytoskeleton for trichome cell morphogenesis (Tian et al., 2015). Furthermore, during mitosis, AtKCBP is found at the cortical division zone, to which the cell plate is eventually guided in the telophase (Buschmann et al., 2015). This localisation was not detected for moss KCBP in the apical cell (Miki et al., 2014). Finally, it has not been verified that walking ability is necessary for KCBP function in organelle transport. An alternative possibility is that KCBP acts as a MT tether, while other motor(s) or MT pushing/pulling force exerted at the cortex drives nuclear and chloroplast translocation. Indeed, MT pushing/pulling is the mechanism of nuclear motility in fission and budding yeasts (Adames and Cooper, 2000; Tran et al., 2001).

In this study, we addressed these questions through a structure-function analysis of KCBP. We identified the FERM-C domain of moss KCBP as critical for cargo transport, which is a dispensable domain of *Arabidopsis* KCBP for MT/actin organisation. In addition, our results suggest that the naked chromatid, rather than the membranous nucleus, binds to KCBP as a cargo in anaphase/telophase, ensuring rapid and robust chromosome segregation to daughter cells.

## Results

### FERM-C domain is required for chromosome and chloroplast translocation

KCBP contains three recognisable domains that are known to bind to other molecules: MyTH4, FERM, and motor domains. To identify the domain(s) responsible for nuclear and chloroplast transport, we made six different truncation constructs of KCBPb, in which one or multiple domains were deleted and Cerulean fluorescent protein was attached (Fig. 1A). We also constructed a control full-length Cerulean-KCBPb and a ‘rigor’ mutant. The rigor mutant possesses a mutation in a residue critical for its motor activity; hence, the motor binds to but does not walk along MTs. The constructs were transformed into a moss line that expresses GFP-tubulin and histoneH2B-mRFP and has had all of its KCBP paralogues deleted, and stable transgenic lines were selected. The expression of each truncated protein was confirmed by immunoblotting (Fig. 1B).

**Figure 1.**
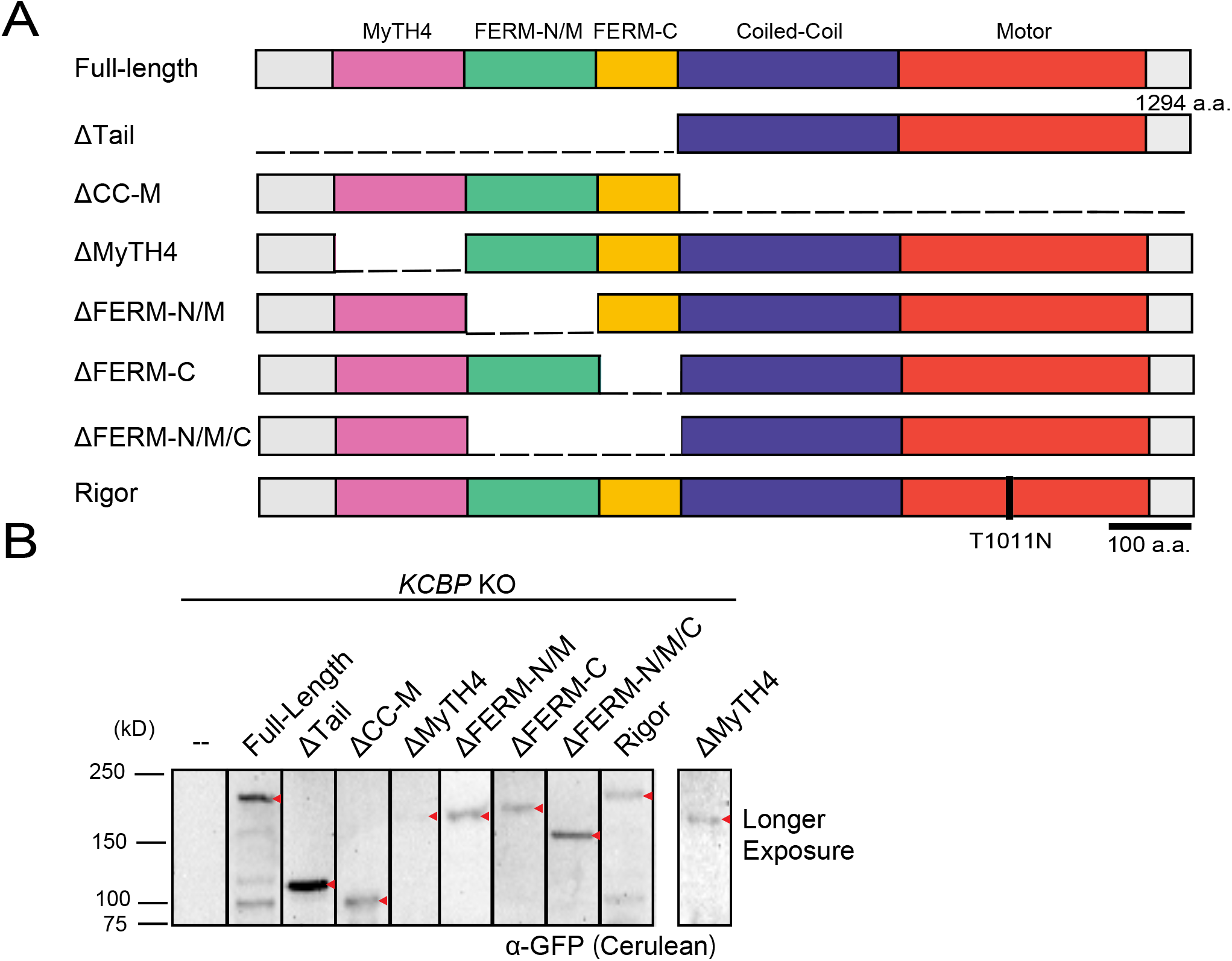
Construction of moss lines expressing truncated KCBP. (A) The eight KCBP constructs used in this study. (B) Immunoblotting of Cerulean-tagged KCBP truncated or mutant constructs by anti-GFP antibody that cross-reacts with Cerulean. The bands of expected size are marked with red arrowheads.

For each truncation, we selected two to four clonal lines and performed long-term time-lapse imaging of the chromosome (histone-RFP) and chloroplast (autofluorescence) with a low magnification lens (Fig. 2A; lines used are #757–764 as listed in Table S1). Applying the previously utilised method (Yamada et al., 2017), we quantified post-mitotic chromosomal/nuclear positioning and chloroplast distribution. As expected, the full-length KCBPb fully restored the positioning of both chromosomes and chloroplasts, validating our assay (Fig. 2B, C). The rigor mutant as well as a motor-less fragment failed to restore the distribution of either organelle, consistent with the model that the positioning of these organelle requires MT-based motility of KCBP (Fig. 2D, E). The FERM domain can be subdivided into three subdomains, FERM-N (also called F1), FERM-M (F2), and FERM-C (F3) (Lu et al., 2014). The fragments that lack FERM-C, which constitutes the lipid- and protein-binding interface, failed to rescue either phenotype, suggesting that this domain is responsible for the organelle attachment of the motor (Fig. 2D, E).

**Figure 2.**
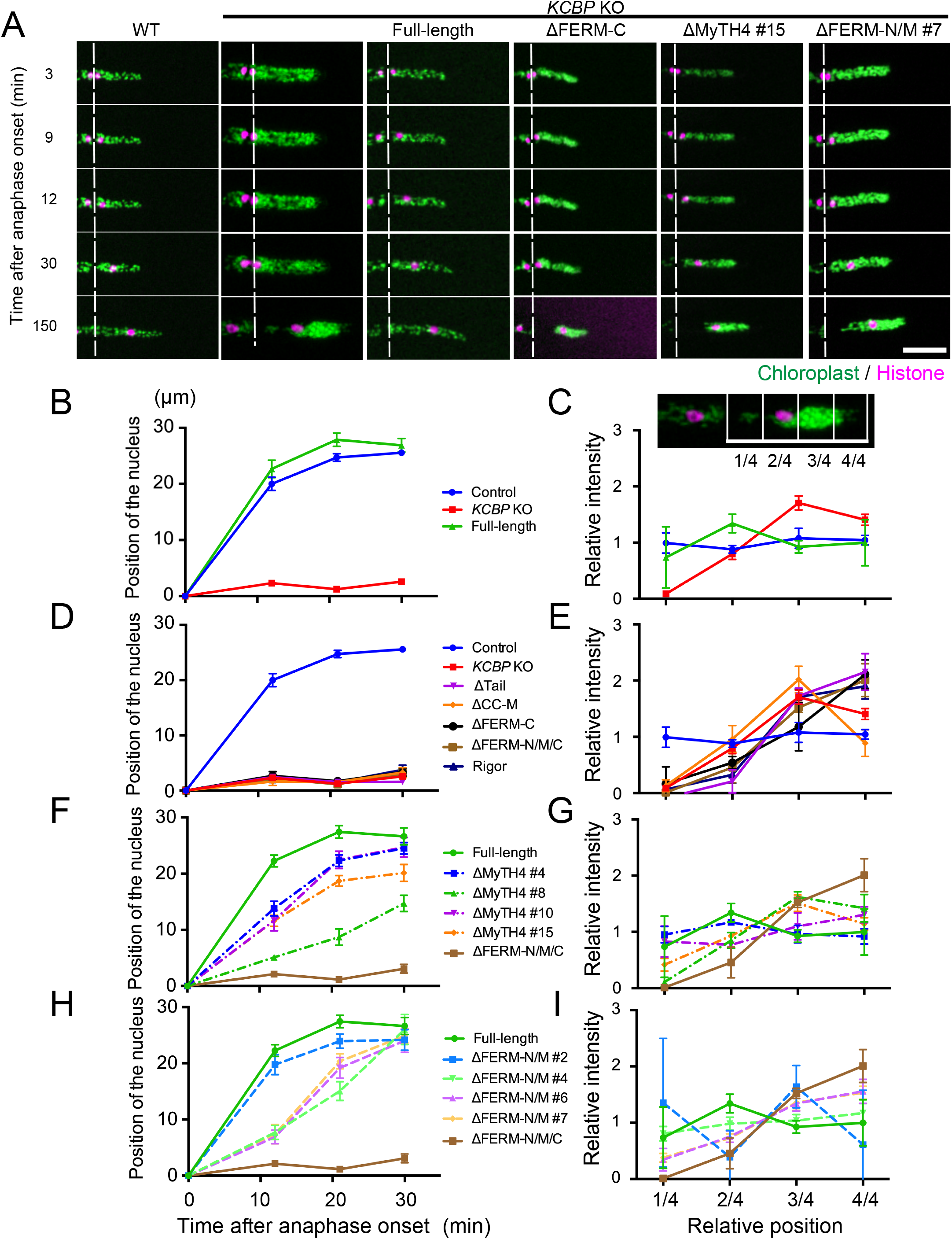
Domains required for chromosome or chloroplast transport. (A) Post-mitotic distribution of the chromosome (histone-RFP, magenta) and chloroplast (autofluorescence, green) in the *KCBP* KO line expressing full-length or 3 truncated constructs. Dashed lines indicate the position of the cell plate. Bar, 50 μm. (B, D, F, H) Position of the chromosome/nucleus after anaphase onset in apical cells. The distance from the metaphase plate is plotted as the mean ± SEM (n = 5). Time 0 corresponds to anaphase onset and the images at 12, 21, and 30 min were analysed. Results from multiple independent lines are displayed for MyTH4 (F) and FERM-N (H) lines, since line-to-line variation was observed. Note that SEM was too small to visualize in some plots. (C, E, G, I) Chloroplast distribution at 150 min after anaphase onset (i.e. during interphase). Following the method described in Yamada et al. (2017), chloroplast signal intensity was measured along cells’ long axes. Each cell was divided into four regions, and the mean signal intensity within each region was divided by the average intensity of the whole cell (schematic illustration is displayed in C). The mean relative values and SEMs are shown in the graph (n = 5). Results from multiple independent lines are displayed for MyTH4 (G) and FERM-N (I) lines, since line-to-line variation was observed.

### MyTH4 and FERM-N/M are not essential for chromosome transport

Specimens lacking either FERM-N/M or MyTH4 domain showed line-to-line variation. In two out of four lines, we observed the rescue of both chromosome and chloroplast translocation, although the chromosome migration rate was somewhat decreased (Fig. 2F–I; ΔMyTH4 #4, #10 lines, ΔFERM-N/M #2, #4 lines). In contrast, in the other two lines, the chloroplast positioning phenotype was not rescued (ΔMyTH4 #8, #15 lines, ΔFERM-N/M #6, #7 lines). The basis of this line-to-line variation is uncertain. Regardless, we observed individual cells that could transport the chromosomes properly but failed to redistribute chloroplasts, indicating that, even when the chloroplast positioning is defective, the chromosomes/nucleus could migrate to its proper position after cell division (Fig. 2A; shown are ΔMyTH4 #15 and ΔFERM-N/M #7 lines).

### KCBP localisation around the chromosomes precedes the completion of NE formation

To correlate the function and localisation of KCBP, we observed chromosomal dynamics as well as mutant KCBP localisation with high-resolution confocal microscopy (Fig. 3). Since our microscope setting could not prevent GFP signal leakage into the Cerulean channel, background GFP-tubulin signal was always detectable when Cerulean-KCBPb fragments were imaged (Fig. 3; ‘Control’ shows a cell that does not express Cerulean). Nevertheless, we were able to observe the chromosomal enrichment of full-length Cerulean-KCBPb during telophase (Fig. 3 ‘Full-length’). The motor-deleted fragment also showed signals on stalled chromosomes, indicating that the cargo-targeting activity was intact in the absence of the motor (‘ΔCC-Motor’). In contrast, ΔTail or ΔFERM-C proteins did not accumulate on the chromosome, decorating phragmoplast MTs instead. The semi-functional MyTH4- or FERM-N/M-deleted fragment showed fainter signals on the chromosomes. These results suggest that accumulation of KCBP is required for chromosome/nuclear transport.

**Figure 3.**
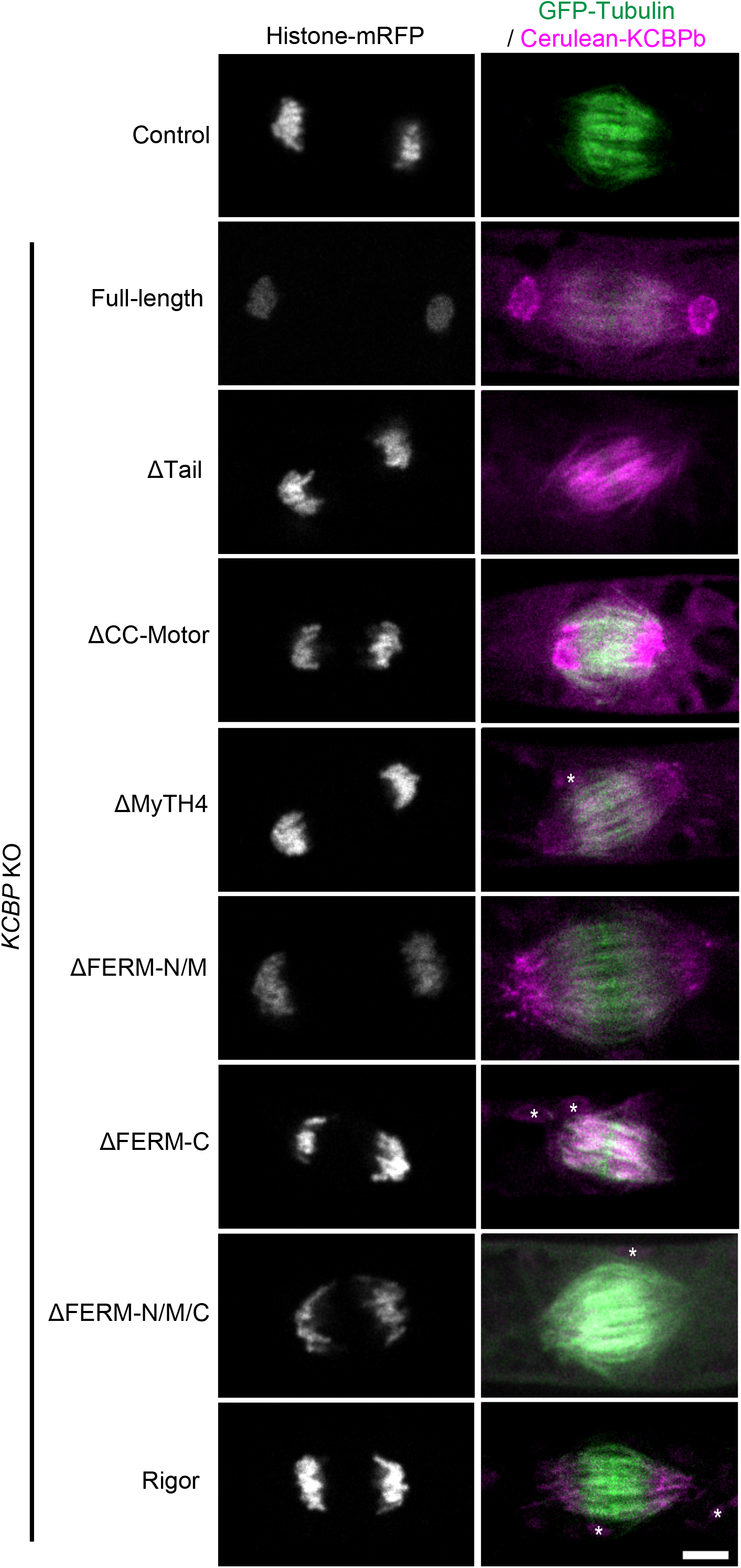
Localisation of truncated KCBP proteins at telophase. Localisation of Cerulean-tagged KCBPb mutants. Presented images are telophase cells (4 min after anaphase onset). Control cells do not possess Cerulean constructs. GFP-tubulin signals partially leaked into Cerulean. Asterisks indicate chloroplast autofluorescence. Bar, 5 μm.

In other cell types, nuclear migration is directed through binding of the motors (kinesin, myosin, dynein) to the NE-inserted protein complex LINC (Chang et al., 2015; Gundersen and Worman, 2013; Tamura et al., 2013). To investigate if this could be the scenario for moss KCBP, we first observed endogenous KCBPb localisation alongside mCherry-tagged SUN1 protein; SUN is a component of LINC and represents NE during re-formation in telophase (Oda and Fukuda, 2011). Interestingly, KCBP accumulation precedes SUN1: SUN1-mCherry signal was not detected around the chromosome at the time of KCBPb-Citrine appearance (Fig. 4A, yellow vs. magenta arrows, Movie 1). Similarly, KCBPb-Citrine appearance precedes that of CRWNa-mCherry, a putative nuclear lamina protein in plants (Fig. 4B–D, yellow vs. magenta arrows, Movie 2) (Pradillo et al., 2019; Wang et al., 2013). These results disfavour the model that KCBP binds to the nucleus via LINC and put into question the assumption that KCBP directly binds to lipid membranes for transport.

**Figure 4.**
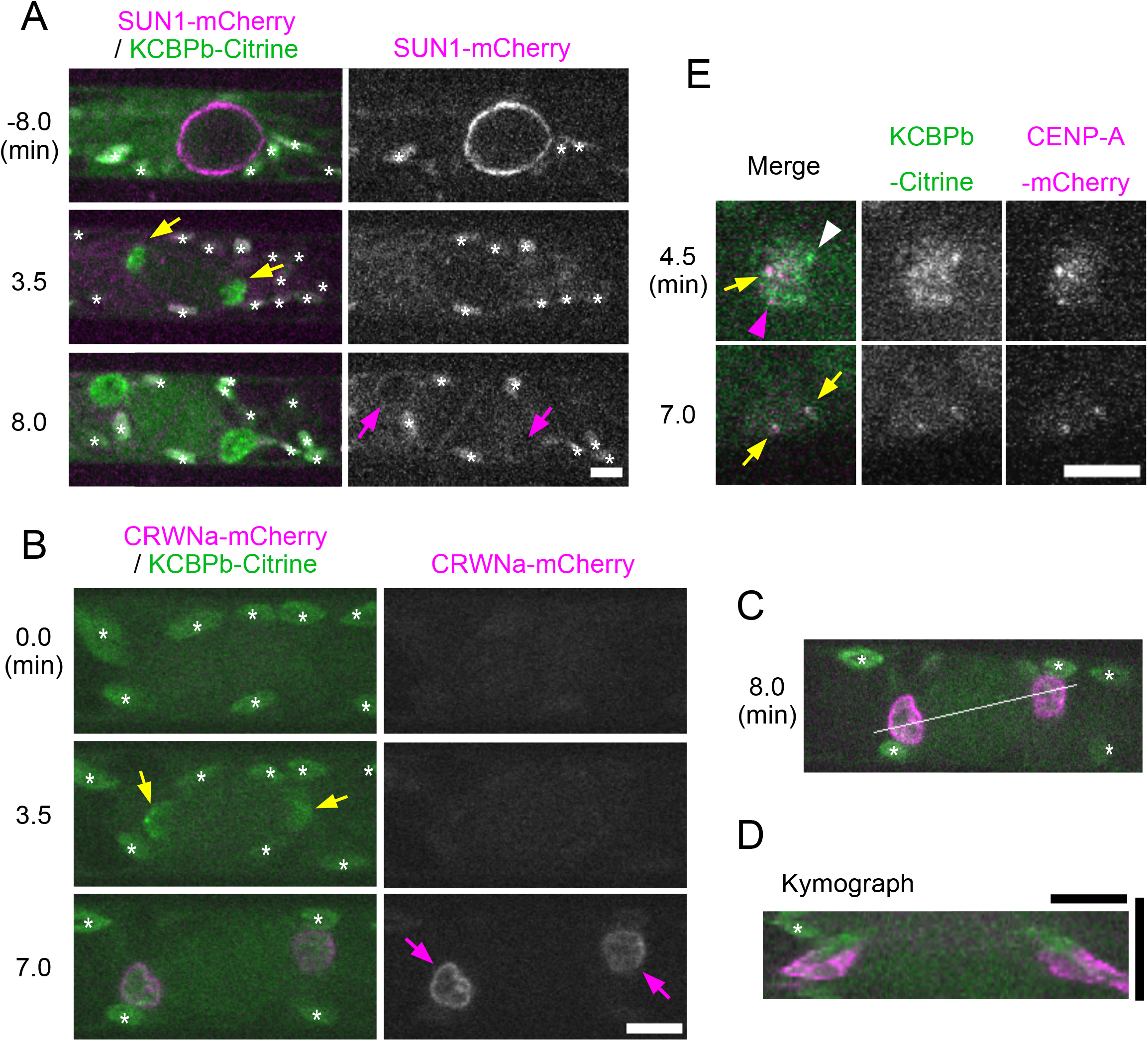
Chromosomal localisation of KCBP precedes completion of nuclear envelope re-formation. (A–D) Chromosomal accumulation of KCBP (yellow arrows) precedes NE localisation of SUN1 or CRWNa (magenta arrows) during telophase. Kymograph of KCBP and CRWNa in (D) was generated based on the white line shown in (C). Bars, 5 μm (A, B, D [horizontal]) and 15 min (D [vertical]). (E) Some punctate signals of KCBP are closely localised with the centromere marker CENP-A-mCherry. Yellow arrows indicate co-localisation, whereas white and magenta arrowheads show non-overlapping signals of KCBPb or CENP-A, respectively. Bar, 5 μm. Asterisks indicate autofluorescent chloroplasts.

Citrine-tagged KCBP showed punctate signals on the chromosomes when imaged with high-resolution microscopy (Yamada et al., 2017). Interestingly, co-imaging of KCBPb-Citrine and CENP-A-mCherry, a constitutive centromeric protein in this cell type (Kozgunova et al., 2019), indicated that some, though not all, punctate signals of KCBP are closely localised with CENP-A (Fig. 4E, yellow arrows and white arrowheads). Conversely, not all centromeres recruited KCBP (magenta arrowheads).

### KCBP accelerates chromatid motility during anaphase

To carefully assess the contribution of KCBP to chromosome segregation in anaphase, we performed time-lapse imaging of *KCBP* KO cells at shorter (10 s) intervals from metaphase (Fig. 5A). Consistent with other imaging results, sister chromatid separation and segregation took place in all KO cells. However, chromosome lagging was detected in 4 out of 7 cells (Fig. 5B; never in 7 control cells) and the rate of chromosome motility, which was quantified with kymographs, was significantly reduced (Fig. 5C–E). This phenotype was not rescued by the expression of ΔFERM-C proteins (3 out of 9 cells showed lagging), whereas the expression of full-length KCBP accelerated chromosome motility relative to the control line (Fig. 5E).

**Figure 5.**
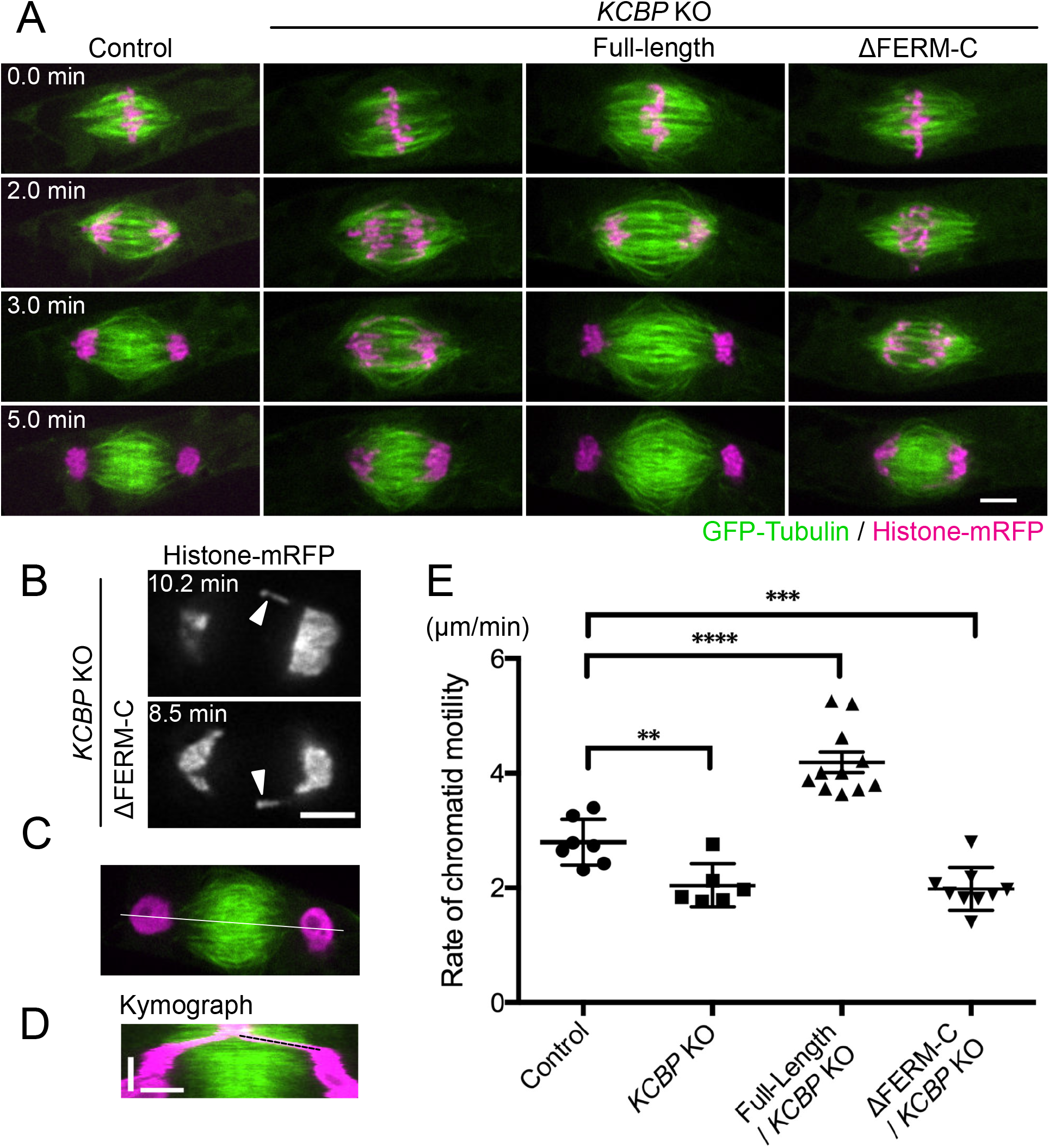
Rate of chromatid motility is reduced in the *KCBP* KO line. (A) Sister chromatid separation and segregation during anaphase. Images were acquired every 10 s. (B) Lagging chromosomes are frequently observed in the absence of FERM-C domain. (C–E) Quantification of the rate of chromosome motility during anaphase based on the kymograph (for example, a kymograph in (D) was generated along the white line shown in (C), and the rate was determined by drawing a dotted line in (D)). Rate of chromatid movement; control, 2.8 ± 0.15 μm/min (±SEM, n = 7); *KCBP* KO, 2.0 ± 0.15 μm/min (n = 6); KCBP full-length add-back, 4.2± 0.17 μm/min (n = 11); ΔFERM-C add-back, 2.0 ± 0.12 μm/min (n = 9). **p < 0.01, ***p < 0.001, ****p < 0.0001 (unpaired t-test, two-tailed). Horizontal bars, 5 μm; Vertical bar in D, 5 min.

### Liverwort KCBP can substitute moss KCBP

To test if the function of KCBP is conserved across plant species, we tried to express *Arabidopsis* and liverwort (*Marchantia polymorpha*) KCBP proteins tagged with Cerulean in the *P. patens KCBP* KO line. After three attempts of transformation, we could not obtain a transgenic line expressing AtKCBP: immunoblotting analysis suggested that the protein is subjected to cleavage and/or degradation (<100 kD band was detected, whereas Cerulean-AtKCBP should be >150 kD; data not shown). We gave up pursuing the functional replacement experiment for AtKCBP. In contrast, we obtained a clonal line in which Citrine-MpKCBP (full-length) expression was confirmed (Fig. 6A). Time-lapse imaging followed by image analysis showed that MpKCBP almost completely suppresses the aberrant positioning of the chromosome and chloroplast (Fig. 6B, C). Furthermore, it showed chromosomal accumulation at telophase (Fig. 6D). Thus, the transport function of KCBP is not endowed specifically to moss.

**Figure 6.**
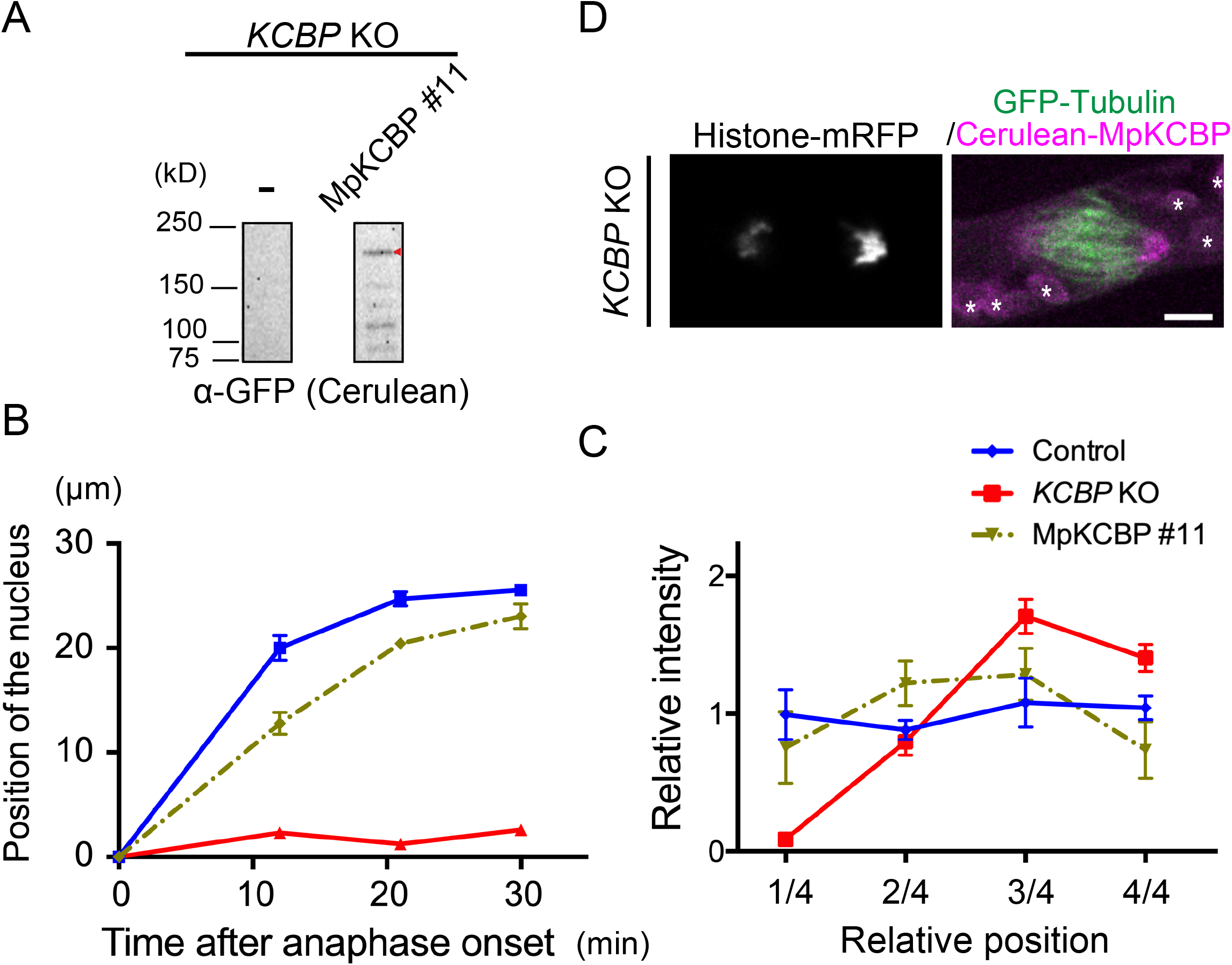
Liverwort KCBP rescues the moss *KCBP* KO phenotype. (A) Expression of Cerulean-tagged *Marchantia polymorpha* (Mp) KCBP in the *P. patens* KCBP KO line. Red arrowhead indicates the expected band of Cerulean-MpKCBP (no Cerulean expression in the left lane). (B, C) Chromosome/nucleus migration (B) and chloroplast distribution (C) were restored by MpKCBP expression in the *PpKCBP* KO line. (D) Nuclear localisation of MpKCBP during telophase. Asterisks indicate autofluorescent chloroplasts. Note that the chromosome on the left was slightly out-of-focus, such that the MpKCBP signal was dimmer than that on the right chromosome. Bar, 5 μm.

## Discussion

### Naked anaphase/telophase chromatids, not the enveloped nucleus, are transported by KCBP

Our previous study showed that, *in vitro*, KCBP binds to phospholipids in a manner dependent on its tail region (Yamada et al., 2017). This led to the model that KCBP might transport membranous organelle, such as nuclei and chloroplasts, through direct binding to their membranes. Although this is still a viable model, our current study urges its reconsideration, at least for the nucleus. We observed that KCBP accumulates on migrating chromosomes before the completion of NE reformation, as indicated by the absence of CRWNa and SUN1. The accumulation of KCBP, particularly near the centromeres, is also not readily explained if KCBP directly binds to NE. The result essentially rejects the hypothesis that LINC acts as the nuclear adaptor for the KCBP motor, unlike in other species. Furthermore, the rate of poleward motility of chromatids reduced and lagging chromatids sometimes appeared during anaphase in the absence of KCBP.

Thus, KCBP likely binds to chromosomal protein(s) at anaphase/telophase and transports chromosomes that are not yet enveloped by lipid membranes (Fig. 7). The importance of the FERM-C domain is not inconsistent with this model, since FERM-C can bind to proteins as well as lipids. In a well-studied case, the crystal structure of the MyTH4-FERM domain attached to a cargo protein was determined for myosins (Hirano et al., 2011; Wei et al., 2011; Wu et al., 2011). Intriguingly, the cargo peptides mainly bind to FERM-N and FERM-C of myosin VII and myoxin X, respectively. In the former, the MyTH4 domain also associates with a part of the cargo protein, likely ensuring its tight association to myosin (Lu et al., 2014; Wu et al., 2011). The MyTH4 domain of KCBP may have a similar function, as MyTH4 deletion from KCBP affects KCBP concentration on chromosomes and the chromosome motility rate. However, the possibility that KCBP might bind to membranes that are being recruited onto the chromosome during NE re-formation cannot be excluded entirely.

**Figure 7.**
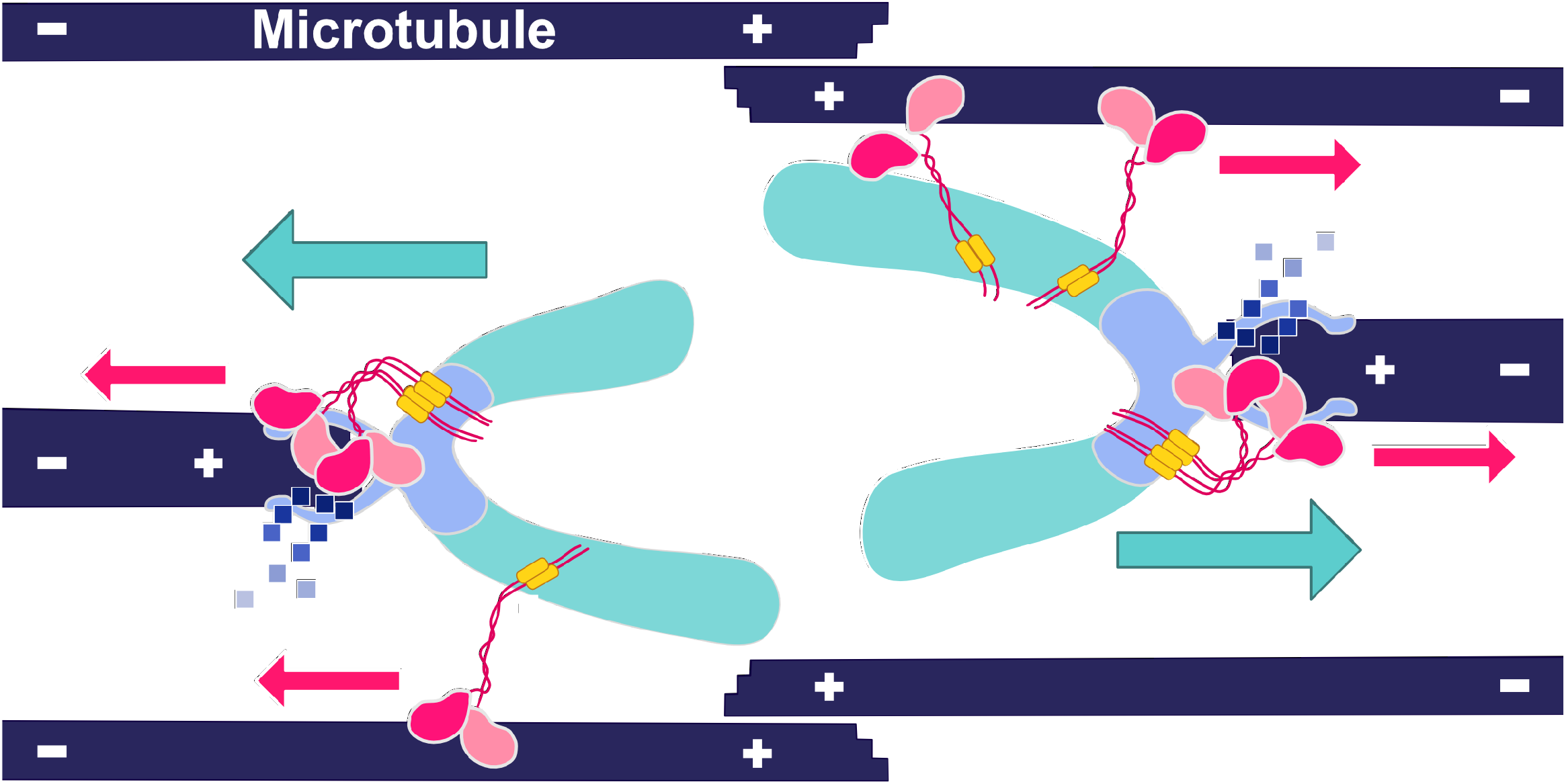
Model for KCBP-dependent transport of chromosomes during anaphase. Minus-end-directed kinesin KCBP (red) binds to the chromatid at the centromere (blue) and other regions (light blue) via the FERM-C domain (yellow) during anaphase. KCBP transports the chromatid as cargo towards the pole, which, together with the depolymerisation of kinetochore-bound MTs (blue boxes; depolymerised tubulin), ensures robust and rapid chromosome segregation into daughter cells.

In contrast to the nucleus/chromosomes, we could not gain much insight into the molecular mechanism of chloroplast transport in this study. The major obstacle was that we could not observe the localisation of KCBP on the chloroplasts, since the autofluorescence of this organelle was overwhelming. However, the essentiality of FERM-C and motor activity and the importance of MyTH4 suggest that KCBP recognises the proteins and/or lipid surface of chloroplasts and transports them to the MT minus-ends, similar to anaphase/telophase chromosomes. Identifying the interacting partner of KCBP at the chloroplast surface could be an interesting future research avenue.

### Conservation of KCBP function

Liverwort KCBP could substitute moss KCBP’s function, suggesting that the FERM-dependent cargo transport activity is preserved in KCBPs of bryophytes. Unfortunately, we could not conduct the complementation analysis for *Arabidopsis* KCBP. However, *Arabidopsis* and moss KCBPs have several common biochemical and structural features. For example, the processive motility of KCBP clusters along MTs has been observed for both *Arabidopsis* and moss KCBPs (Jonsson et al., 2015; Tian et al., 2015). The domain arrangement in the tail is well conserved; in particular, the FERM-C domain (~108 a.a.) is highly conserved (93 % identity in amino acid sequences). Nevertheless, previous study showed that the entire MyTH4-FERM region is dispensable for KCBP function in *Arabidopsis* trichome morphogenesis (Tian et al., 2015). Thus, the two KCBP functions – the FERM-dependent transport and the FERM-independent MT-actin organisation – are separate. We speculate that AtKCBP also carries out the FERM-dependent transport function, possibly redundantly with other motors, in certain tissues of *Arabidopsis*.

Poleward chromatid motility during anaphase A is mainly driven by depolymerisation of kinetochore-bound MTs (Asbury, 2017). However, it was reported in flies that a kinetochore-enriched dynein motor accelerates the motility of chromatids and prevents their lagging, perhaps by transporting the chromatid as cargo (Savoian et al., 2000; Sharp et al., 2000). This is highly analogous to what we observed for KCBP in the present study (Fig. 7). More recently, it was shown that another subclass of kinesin-14 in maize localises to and moves neocentromeres along spindles to promote meiotic drive (Dawe et al., 2018). Thus, chromatid transport in anaphase A by minus-end-directed motors may be a widely utilised strategy to ensure robust transmission of genetic materials. In moss, this role is fulfilled by KCBP.

## Materials and methods

### Moss culture and transformation

Moss lines used in this study are listed in Table S1; all lines originated from the *Physcomitrella patens* Gransden2004 strain. Methodologies of moss culture, transformation, and transgenic line selection have been thoroughly described in previous studies (Yamada et al., 2016). Briefly, cells were cultured on BCD agar medium for imaging. Transformation was performed by the standard PEG-mediated method and stable lines were selected with antibiotics. The *mCherry* gene was inserted into the C-termini of *CRWNa* and *SUN1* via homologous recombination. Citrine and mCherry tag insertion were confirmed by PCR. Integration of Cerulean-KCBP truncation constructs was confirmed by PCR followed by immunoblotting.

### Plasmid construction

Plasmids and primers for plasmid construction used in this study are listed in Table S2. To generate truncation/rescue plasmids, *KCBP* sequences were amplified from a cDNA library and ligated into the pENTR/D-TOPO vector containing *Cerulean* sequences, followed by a Gateway LR reaction into a pMN601 vector that contains the EF1α promoter, nourseothricin-resistance cassette, and *PTA1* sequences designated for homologous recombination-based integration (Miki et al., 2016). *Arabidopsis* and *Marchantia KCBP* sequences were similarly cloned into a pTM153 vector that contains the EF1α promoter, blasticidin-resistance cassette, and *PTA1* sequences designated for homologous recombination-based integration (Miki et al., 2016). For endogenous CRWNa or SUN1 tagging with mCherry, a plasmid was constructed, where the *mCherry* gene and the blasticidin-resistance cassette were flanked by ~1 kb sequences within the ORF and 3′UTR. CENP-A-mCherry driven by actin promoter was targeted to hb7 locus and selected by nourseothricin resistance.

### Immunoblotting

Cell extracts were prepared by grinding protonema colonies (Yamada et al., 2016). Immunoblotting of Cerulean-tagged proteins was performed with home-made anti-GFP antibody (rabbit “Nishi”, final bleed, 1: 500).

### In vivo microscopy

Methods for epifluorescence and spinning-disc confocal microscopy were previously described (Yamada et al., 2016). Briefly, protonemal cells were plated onto glass-bottom plates coated with BCD agar medium and cultured for 6–7 d. Long-term imaging by a wide-field microscope (low magnification lens) was performed with a Nikon Ti (10× 0.45 NA lens and EMCCD camera Evolve [Roper], controlled by Micromanager software) or a Nikon TE (10× 0.45 NA lens and CMOS Camera ZYLA-4.2P-USB3 [Andor], controlled by iQ software). High-resolution imaging of Citrine and mCherry (or mRFP) was performed with a spinning-disc confocal microscope (Nikon Ti; 100× 1.45 NA lens, CSU-X1 [Yokogawa], and EMCCD camera ImagEM [Hamamatsu]). Cerulean imaging was performed with another spinning-disc confocal microscopy (CSU-W1 was attached to IX-83: Olympus) that was controlled by Metamorph software. All imaging was performed at 24–25°C in the dark.

### Data analysis

Quantification of the chromosome and chloroplast movement following cell division was performed for apical cells, following the method described in Yamada et al. (2017). Images were acquired every 3 min for 10 h under dark condition and kymographs were generated. The distance from the metaphase plate to the nuclear centre was measured after generating kymographs. To quantify chloroplast distribution, the intensity of chloroplast autofluorescence was measured. Images were acquired every 3 min for 10 h under dark condition. Chloroplast signal intensity was measured along cells’ long axes by drawing a one-pixel-wide line using ImageJ. Each cell was divided into four regions, and the mean signal intensity within each region was divided by the average intensity of the whole cell.

### Accession number

*Physcomitrella patens SUN1*; Pp3c7_4170: *CRWNa*; Pp3c1_1360: *Marchantia polymorpha KCBP*; Mapoly0012s0174.

## Acknowledgements

We thank Elena Kozgunova for CENP-A-mCherry plasmid; Rie Inaba for media preparation; Yoshikatsu Sato and Advanced Imaging Support (ABiS) platform (JP16H06280) for help of confocal imaging; and Elsa Tungadi for manuscript editing. This work was funded by JSPS KAKENHI (17H06471, 18KK0202) and by JSPS and DFG under the Joint Research Projects-LEAD with UKRI to G.G. M.Y. was a recipient of a JSPS pre-doctoral fellowship (16J02796).

## Figure and movie legends

**Movie 1. Co-imaging of KCBPb-Citrine and SUN1-mCherry during telophase**

Images were acquired every 30 s with spinning-disc confocal microscopy. Arrowheads indicate Citrine/mCherry signals, whereas many other signals represent chloroplast autofluorescence. Bar, 5 μm.

**Movie 2. Co-imaging of KCBPb-Citrine and CRWNa-mCherry during telophase**

Images were acquired every 30 s with spinning-disc confocal microscopy. Arrowheads indicate Citrine/mCherry signals, whereas many other signals in the Citrine channel represent chloroplast autofluorescence. Bar, 5 μm.

**Movie 3. Reduction in the rate of chromatid motility in anaphase in the *KCBP KO line***

Images were acquired every 10 s with spinning-disc confocal microscopy. Arrowheads indicate lagging chromosomes. Green, GFP-tubulin; Magenta: histoneH2B-mRFP. Bar, 5 μm.

**Table S1.**
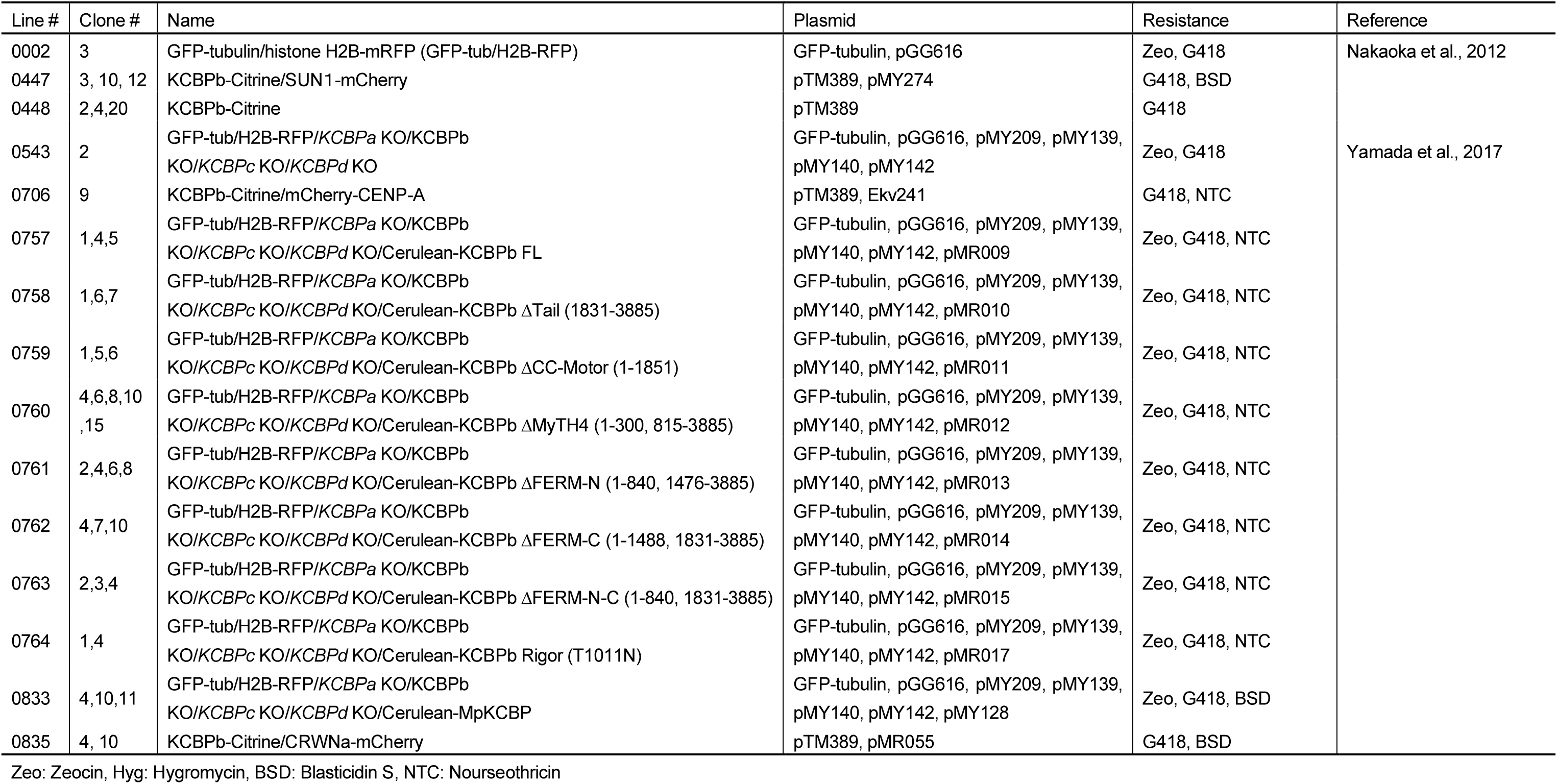
Transgenic moss lines used in this study.

**Table S2.**
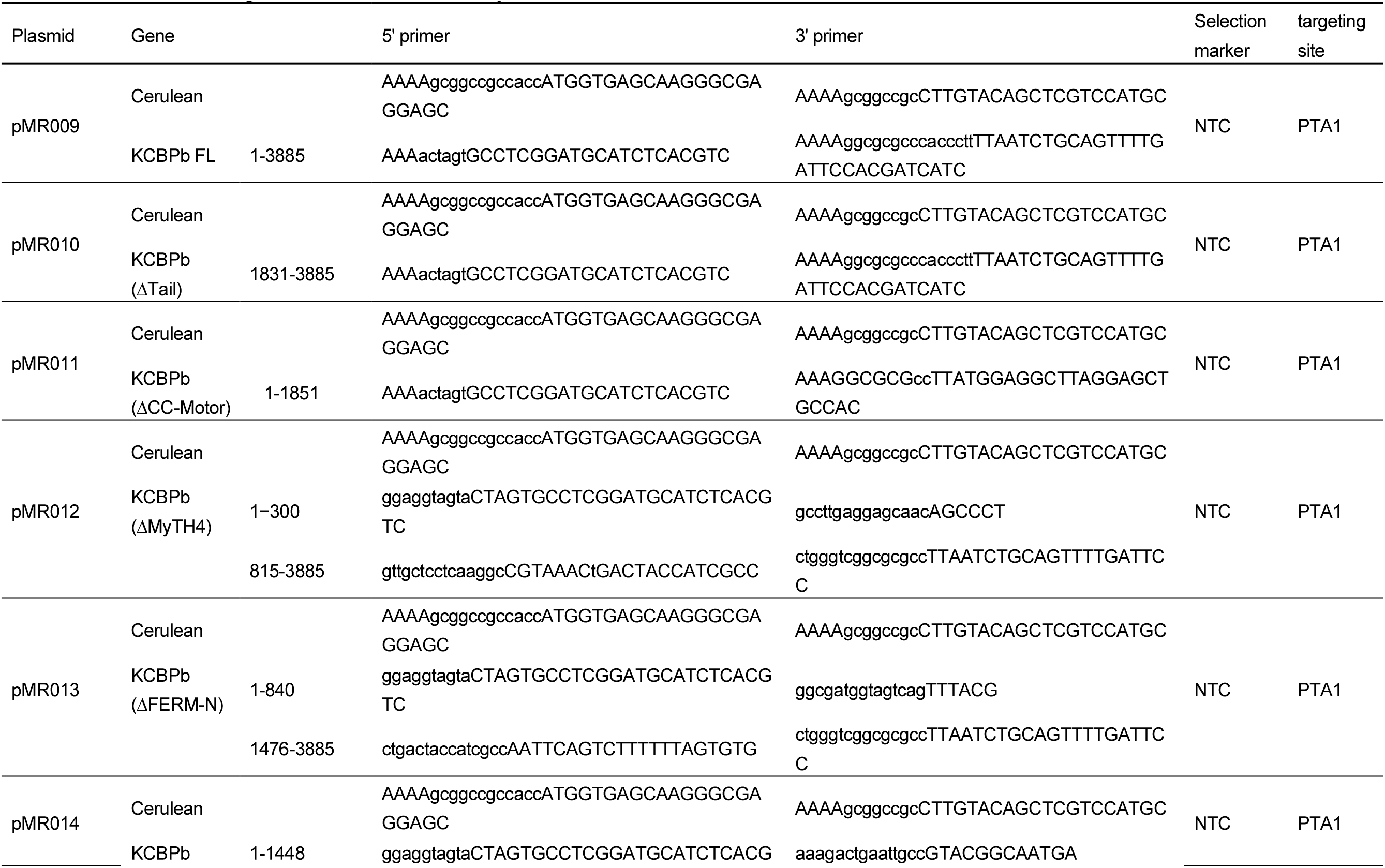
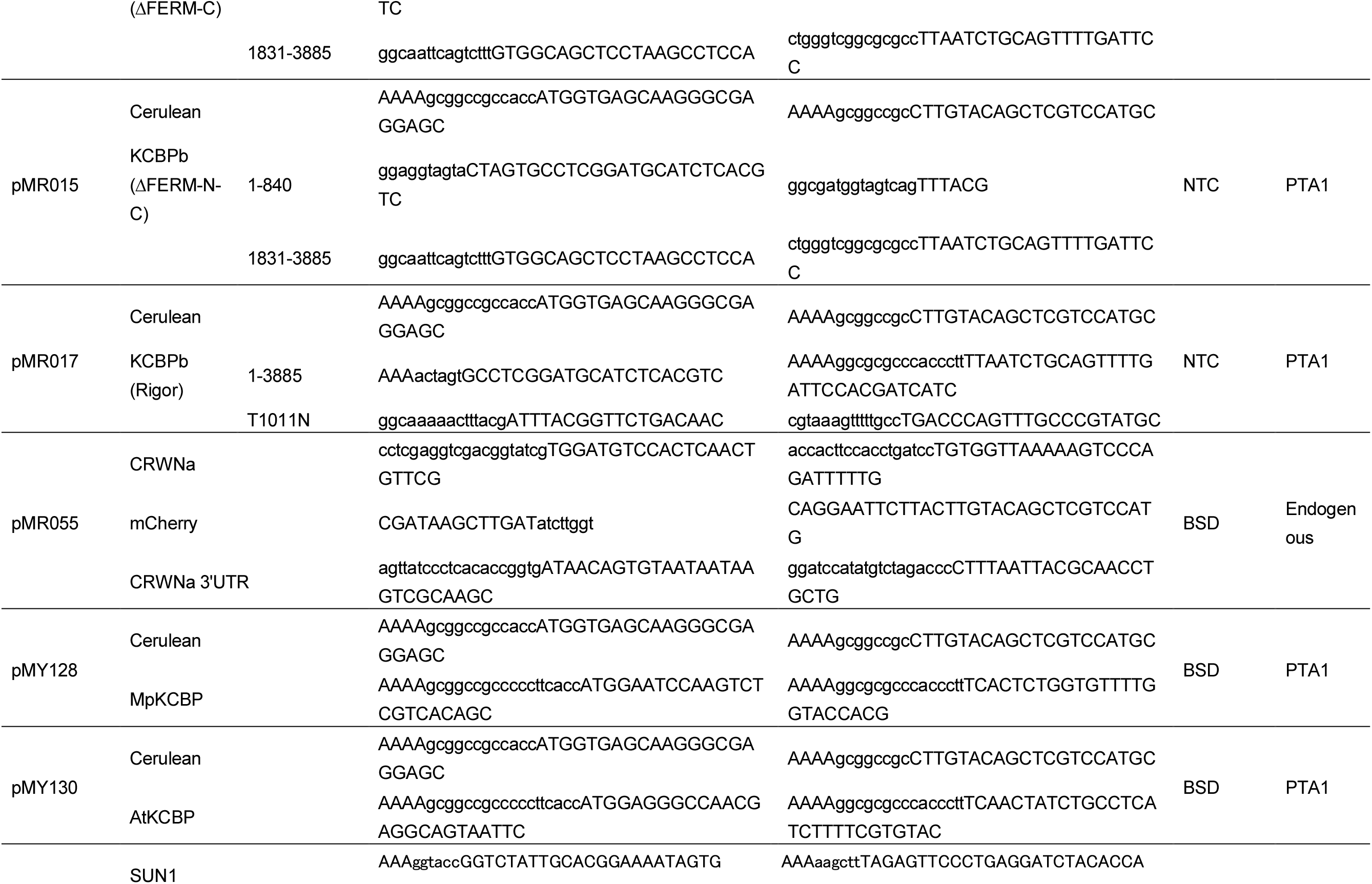
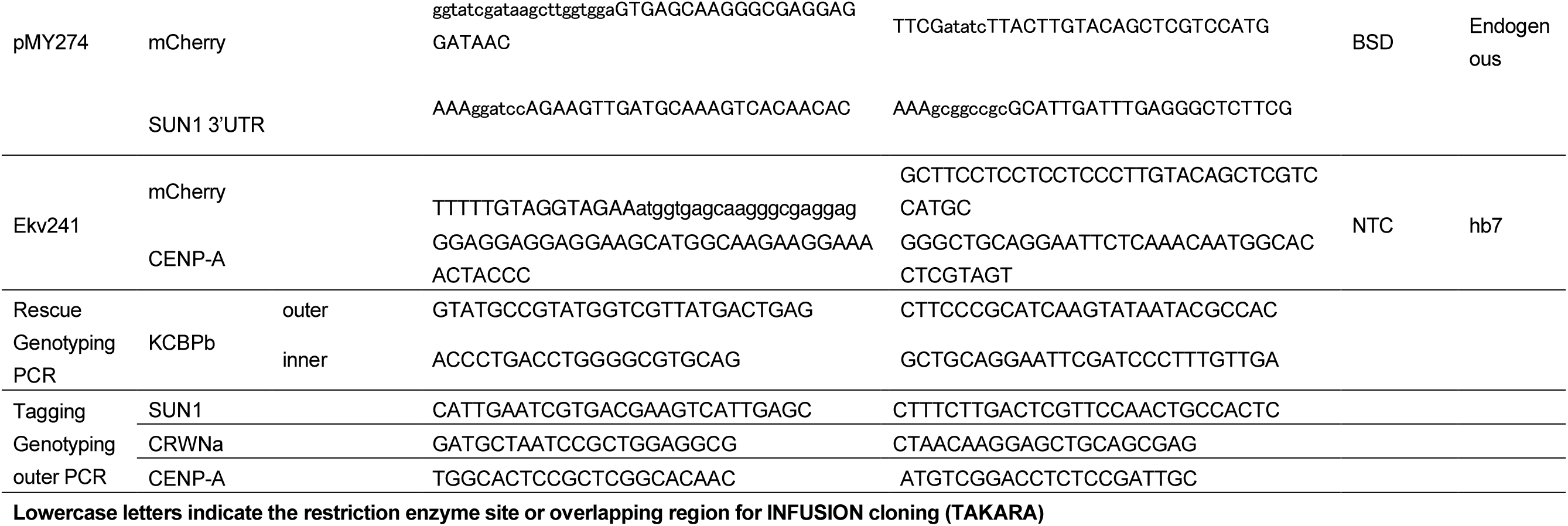
Plasmids and primers used in this study.

